# Differential Equation Modeling of Cell Population Dynamics in Skeletal Muscle Regeneration from Single-Cell Transcriptomic Data

**DOI:** 10.1101/2024.11.06.622368

**Authors:** Renad Al-Ghazawi, Xiaojian Shao, Theodore J. Perkins

## Abstract

Skeletal muscle regeneration is a complex process orchestrated by diverse cell populations within a dynamic niche. In response to muscle damage and intercellular signaling, these cells undergo cell fate and migration decisions including quiescence, activation, proliferation, differentiation, infiltration, apoptosis, and exfiltration. The emergence of single-cell RNA sequencing (scRNA-seq) studies of muscle regeneration offers a significant opportunity to refine models of regeneration and enhance our understanding of cellular interactions. To better understand how crosstalk between cell types governs cell fate decisions and cell population dynamics, we developed a novel non-linear ordinary differential equation model guided by scRNA-seq data. Our model consists of 9 variables and 19 parameters, capturing the dynamics of key myogenic lineage and immune cell types. We calibrated time-series scRNA-seq data to units of cells per cubic millimeter of tissue and fit our model’s parameters to capture the observed dynamics, validating on an independent time series. The model successfully captures regeneration dynamics, particularly after incorporating a novel type of regulatory interaction between M2 macrophages and satellite cells that has been hypothesized in the literature. Our model lays a foundation for future computational explorations of muscle regeneration, modeling of disease conditions, and in silico testing of therapeutic strategies.

## 1 Introduction

Skeletal muscle function relies on the coordinated activity of multinucleated myofibers, each housing hundreds to thousands of nuclei (myonuclei) located at the periphery under the sarcolemma [40]. These myonuclei originate from muscle stem cells, also called muscle satellite cells (MuSCs), which form a heterogeneous population with distinct states and functions [38, 52, 56]. Under homeostatic conditions, MuSCs reside in a quiescent state characterized by Pax7 expression and low metabolic activity, ensuring their longevity and readiness for tissue regeneration [22]. This pool of quiescent MuSCs (QSCs) serves as a vital reservoir for regeneration, and disruptions in their maintenance, such as mutations or chronic damage, can lead to depletion and impaired muscle repair. When exposed to signals from a damaged environment, QSCs become activated satellite cells (ASCs) and display their dynamic nature by responding to various signaling pathways, and by upregulating Myod1 and Myf5, among other genes [2]. These activated cells can then differentiate into myocytes, which later fuse into myofibers, or return to quiescence, replenishing the QSC pool to support future regeneration. Notably, this intricate process is not solely driven by MuSCs and their distinct states, but also involves complex interactions with various cell types within the muscle niche, including immune cells such as neutrophils, monocytes, and macrophages.

Computational modeling of muscle regeneration has been used to formulate and test hypotheses on different aspects of the regeneration process. These models generally use one of two formalisms, differential equations [9, 21, 55, 15, 28, 50] or agent-based simulations [36, 37, 64, 48, 63, 66, 26, 17]. Differential equation-based models are typically concerned with the amounts of different types of cells or myofibers and their fate decisions. These models typically ignore spatial information, but offer rapid simulation and analytical tractability. Agent-based models are typically slower to simulate and less amenable to mathematical analysis, but offer the capability to express more detailed cell fate decision rules, and to explore roles of tissue architecture and cell migration. Because of the interplay between immune cells and cells in the myogenic lineage, many works have focused on those relationships [9, 21, 55, 28, 50]. For instance, Stephenson and Kojouharov [55] modeled the dynamics of macrophages and their influence on myogenic cells, exploring single-and multiple-injury scenarios, and regeneration in aged muscle. Because muscular dystrophy is one of the most common genetic disorders [13], many works have focused on modeling the dysfunction of regeneration in that scenario [10, 21, 26, 64, 63]. For instance, Virgilio et al. [64, 63] explored the role of fibrosis in muscle regeneration, and whether reducing fibrosis might aid regeneration in the dystrophic context. Earlier work by Jarrah et al. [21] focused on the role of immune cell types in muscular dystrophy, predicting substantial roles for and changes in those cells over the course of months as the disease progresses. Farhang-Sardroodi and Wilkie [15] modeled cancer cachexia, rapid muscle-wasting associated with cancer, and its potential for therapeutic treatment by blocking the myostatin/activin A pathway.

Single-cell RNA-sequencing (scRNA-seq) has enabled quantification of gene expression levels across thousands of individual cells within a single controlled experiment [29, 20]. In stem cell biology, scRNA-seq has been used extensively to study differentiation processes (e.g. [5, 42, 16, 53, 51]), and to identify subpopulations of stem cells with distinct features (e.g. [34, 35, 31]). In the context of muscle regeneration biology in particular, there has considerable use of scRNA-seq (see McKellar et al. [39] for a comprehensive integration and analysis of different datasets) and several novel discoveries. For instance, De Micheli et al. [7] identified an important role for Syndecan proteins in mediating stage-specific communications in satellite cells. Oprescu et al. [45] identified a novel subpopulation of MuSCs termed immunomyoblasts (IMBs). Lazure et al. [33] studied transcriptional and chromatin changes in MuSCs in aging mice, showing that those MuSCs can be substantially rejuvenated by exposure to the MuSC niche of young mice.

Despite the many previous computational studies of muscle regeneration, and despite the many scRNA-seq studies of muscle regeneration, we know of no modeling studies that explicitly take advantage of scRNA-seq data in their formulation or parameter fitting. In the present work, we formulate a novel mathematical model of muscle regeneration, focusing on cell fate decisions and cell-cell communication between myogenic and immune cell types. The primary biological question we address is whether known channels of communication between cell types are sufficient to explain observed cell population dynamics from scRNA-seq time-series of standard experimental muscle injury and regeneration protocols. The two primary technical challenges we faced were (1) how to normalize single-cell RNA-seq time-series data to provide absolute, not merely fractional, cell population size estimates; and (2) how to formulate a mathematical model of muscle regeneration dynamics of complexity suitable to the data. We formulate a differential equation model similar to several earlier models (esp. [55, 28, 50]), but different in some details. Our primary observation is that while established cell-cell interactions are largely sufficient to explain observed cell fate decisions and population dynamics, it was necessary to include in the model a link from M2 macrophages to activated satellite cells that promotes deactivation—including both differentiation towards the myocyte fate for some cells, but also a return to quiescence for other cells that renews the satellite cell pool.

## 2 Results

Injury-regeneration protocols are by far the most common means of studying muscle regeneration in vivo, and typically consist of injuring a muscle in a test animal by toxin injection or crushing [8, 19]. The muscle then goes through several phases of regeneration, including: an inflammatory phase, in which immune cells infiltrate the muscle and clean up damage tissue, while MuSCs activate and start to proliferate; an anti-inflammatory phase, in which immune cells exfiltrate and MuSCs proceed to differentiate; and a final return to homeostasis, in which a few MuSCs return to quiescence, while most fuse into mature, functional myofibers.

We begin by describing the structure and equations of our ordinary differential equation (ODE) model of this process, which is based on known cell fate regulatory interactions. We then describe how we used mouse scRNA-seq data, collected under a standard injury-regeneration protocol, to produce calibrated time-series that the model should be able to reproduce. We use nonlinear least squares (NLLS) optimization to optimize model parameters to fit the calibrated time-series data, validating the model on independent injury-regeneration time-series data from a different scRNA-seq study. Lastly, we report sensitivity analysis to understand the influence of each parameter on the model dynamics.

### 2.1 Mathematical formulation of cell population dynamics during skeletal muscle regeneration

The population dynamic model we present was developed from literature and preceding models [21, 69, 11, 32, 58, 44, 41, 1, 60, 59], encapsulating the interactions among nine primary cell types: damaged myonuclei (Md), neutrophils (N), apoptotic neutrophils (Nd), monocytes (M), classically activated macrophages (M1), alternatively activated macrophages (M2), quiescent satellite cells (QSC), activated satellite cells (ASC), and myocytes (Mc) (Fig. 1). It includes both cell fate decisions and cell-cell communication processes that influence those decisions. Each variable represents the number of that cell type present as a function of time after an injury event, in units of cells per cubic millimeter. These cell population numbers are governed by differential equations specified below. Rates of cell activities, such as their infiltration or coming into existence (influx), transforming into another type (differentiation/polarization), apoptosis or leaving (outflux), and affecting each other (signaling), are determined by rate constants denoted by c, as well as numbers of cells performing or regulating the activity. It should be noted that due to the limitations of scRNA-seq in capturing apoptotic neutrophils and damaged myonuclei, empirical data for these variables were not available. Consequently, the model training described below focused on live cell types where empirical data were obtainable (N, M, M1, M2, QSC, ASC, and Mc). The fitting of these variables effectively informed the dynamics of the other model variables, for which empirical data were not directly measured. Below, we explain each equation in detail.

**Figure 1.**
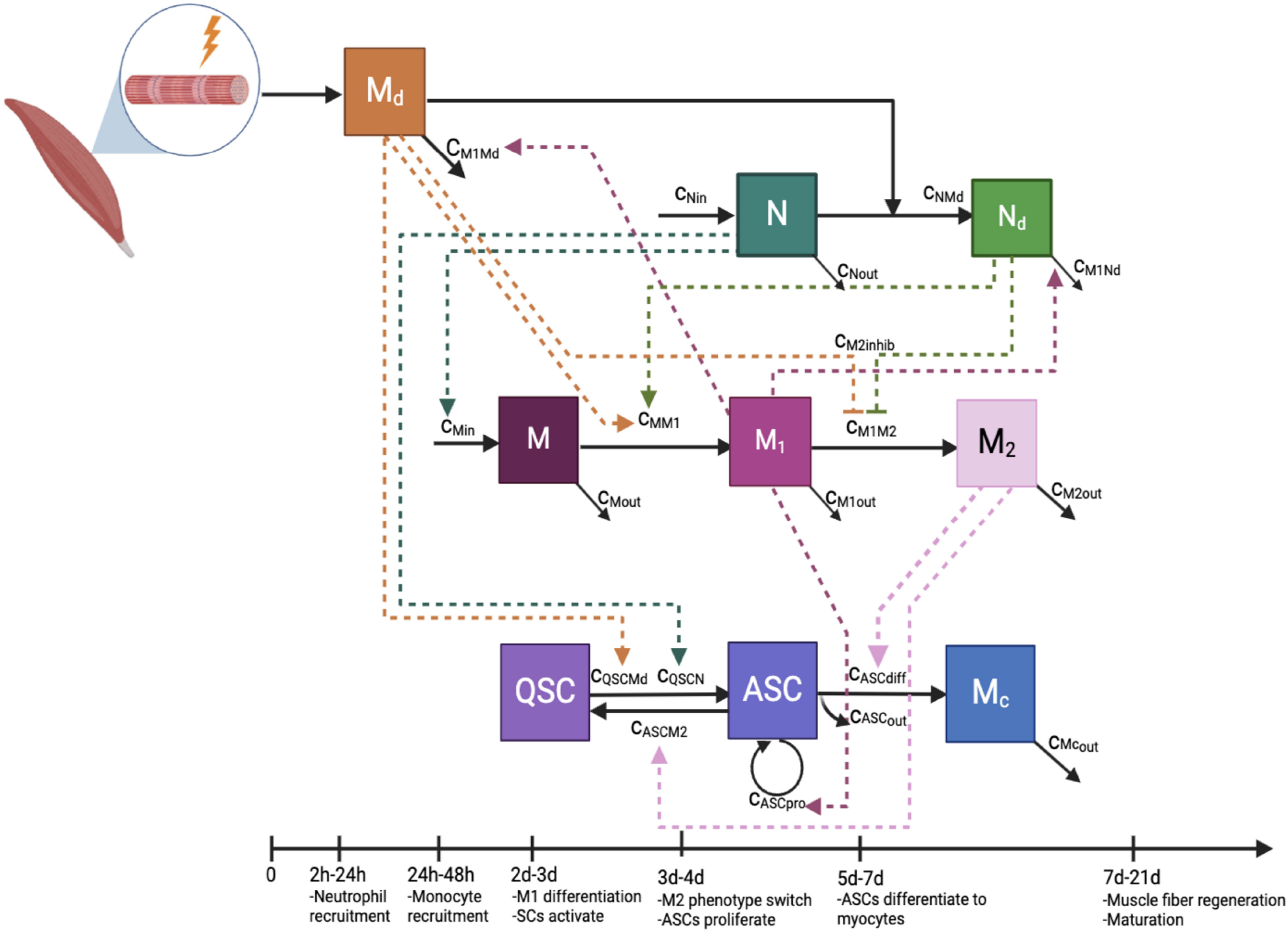
Schematic Representation of Cellular Dynamics in Muscle Regeneration. The diagram delineates the interactions among various cell types involved in the muscle regeneration process following injury, including the sequential recruitment and transformation of cells. Solid arrows represent transformations/transitions between cellular states, while dashed arrows indicate regulatory influences. The estimated times of major events after injury (at day 0) are indicated at the bottom of the diagram. Initially, damaged myonuclei (Md) trigger an inflammatory response. The first responders, neutrophils (N), migrate to the injury site and transition into apoptotic neutrophils (Nd) upon consuming damaged myonuclei, and are subsequently cleared by classically activated macrophages (M1). Monocytes (M) are recruited by neutrophils and rapidly differentiate into M1 macrophages, which later transition into alternatively activated macrophages (M2) as injured tissue is cleared. Quiescent satellite cells (QSC) activate to activated satellite cells (ASC) in response to the damage. Activated satellite cells proliferate until, under the influence of M2 macrophages, they either differentiate and fuse to form myocytes (Mc) or revert to the quiescent state, contributing to the pool of QSCs. Myocytes eventually fuse into multinucleated myofibers, but our model does not include this process.

#### Muscle damage and clearance

Muscle injury is represented by a non-zero initial value of damaged myonuclei, Md. Dead myonuclei are cleared away both by neutrophils and by M1 macrophages. Neither cell type is initially present in the muscle in substantial numbers, but as we will describe below, immune cells are recruited to the damaged muscle as the part of an inflammatory process driven by pro-inflammatory cytokines, growth factors, and metabolites [70, 61, 65]. Once present, they exert their influence in part by clearing the dead myonuclei. Therefore, we model the rate of change of Md (Eq. 1), or equivalently their clearance rate, as depending on the presence of M1 macrophages at the rate of *c*_*M*1*Md*_·*M* 1·*Md* and the presence of neutrophils at the rate of *c*_*NMd*_ · *N* · *Md* .

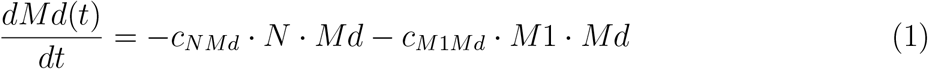

#### Neutrophil recruitment and activity

Neutrophils phagocytose Md and subsequently secrete cytokines that trigger a cascade of cellular responses essential for orchestrating muscle regeneration [68]. To derive the rate of change of N, we consider three processes: the influx of neutrophils into the damaged area, their natural death or exit from the system, and their activity in phagocytosing Md (Eq. 2). The influx correlates proportionally with the level of damage, as indicated by Md. This correlation is supported by empirical evidence showing that inflammation intensity varies with the extent of injury [46, 62, 43]. The linear term 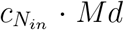 represents the influx of neutrophils and the constant 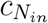 is the rate at which neutrophils are recruited in response to chemotactic signals released by the damaged muscle. A first-order decay process 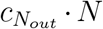 models the natural decay or exit of neutrophils from the system. The bilinear term 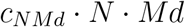 models the neutrophil activity in phagocytosis of Md. This synergistic interaction not only reduces the population of damaged cells but also leads to the apoptosis of neutrophils post-phagocytosis. The apoptotic neutrophils will leave debris that macrophages clear up, which provides a signal for the resolution of inflammation [67]. These three processes combine to generate the neutrophil dynamics equation as follows.

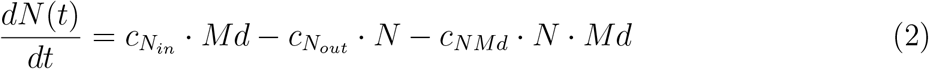

To derive the rate of change for apoptotic neutrophils Nd (Eq. 3), we consider their formation from N through phagocytosis and the clearance by M1 macrophages. This conversion, directly dependent on the phagocytosis activity, is modeled by the bilinear interaction in Eq. 2. Their clearance by M1 macrophages is modeled with a second interaction, dependent on both their population and the population of M1, with *c*_*M*1*Nd*_ · *M* 1 · *Nd* representing the rate of this clearance process.

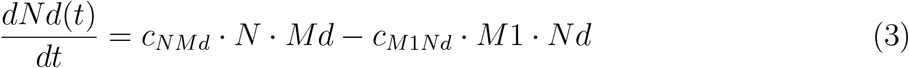

#### Monocyte influx and polarization to M1 macrophages

Following neutrophil recruitment, monocytes are mobilized and their transition to the M1 phenotype occurs typically within 24 hours [47]. The concentration of M1 macrophages is reported to peak approximately two days post-injury (DPI), parallel with the decline of neutrophils, facilitating the clearance of remaining Md and Nd [54]. We consider three processes for monocyte dynamics: their recruitment to the injury site, their transformation into M1 macrophages, and their natural exit from the system (Eq. 4). The recruitment to the site of injury is driven N, assuming a direct, proportional relationship, modeled by the linear term 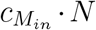 They then polarize into M1 macrophages. This transformation is dependent on both the presence of Nd and/or Md, leading to the term *c*_*MM*1_ · *M* (*Nd* + *Md*) reflecting the influence of apoptotic neutrophils and damaged muscle on monocytes. The natural exit of monocytes from the system is given by 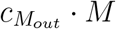.

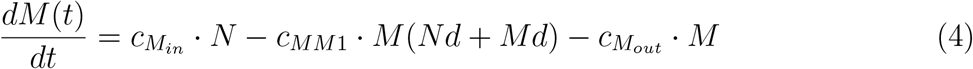

#### Macrophage-mediated regeneration

Macrophages play a biphasic role in regeneration, where M1 macrophages foster the proliferation of ASCs while inhibiting their differentiation. Conversely, M2 macrophages suppress proliferation but encourage differentiation and fusion [1, 70, 23, 12]. Upon completion of phagocytosis and clearance of cellular debris, M1 macrophages undergo a phenotypic switch to alternatively activated M2 macrophages. This transition facilitates the differentiation and fusion of satellite cells.

The dynamics of M1 and M2 macrophages are derived by considering their transformation from monocytes, transition between states, and exit from the system (Eqs. 5 & 6). M begin to differentiate into M1 influenced by signals from dead myonuclei or dead neutrophils, which we model by the term *c*_*MM*1_ · *M* (*Nd* + *Md*). Conversely, the progression from M1 to M2 macrophages is repressed by signals from dead myonuclei or neutrophils represented by the term 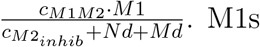that do not convert to M2s subsequently exit the system at a rate of 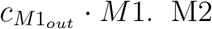 macrophages exert influences on satellite cell behavior (see below) and eventually exit the system at a rate of 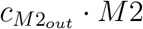. This results in the following equations:

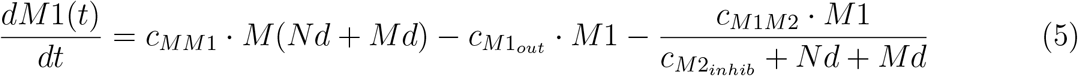

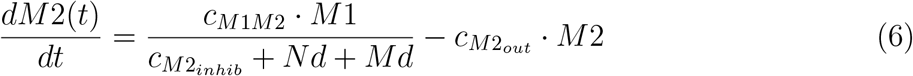

#### Satellite cell dynamics

In the absence of injury, satellite cells maintain a quiescent state within the muscle fiber [22, 69]. However, muscle damage triggers rapid satellite cell proliferation, a response that can occur independently but is amplified by the presence of M1 macrophages [47, 12]. The model also considers that myocytes, derived from ASCs, align and fuse under the influence of M2 macrophages to regenerate healthy muscle fibers [58]. If no further damage occurs, M2 macrophages are assumed to exit the system by either dying or migrating to other regions, reflecting the self-limiting nature of the regenerative response. This assumption aligns with observations that the resolution of inflammation coincides with the cessation of active regeneration and the re-establishment of muscle homeostasis [58, 70]. Therefore, our model incorporates for satellite cells the dynamics of both self-renewal and differentiation processes.

In our QSC dynamics equation (Eq. 7), the first term is bilinear, *c*_*QSCN*_ · *QSC* · *N*, which represents the conversion of QSCs as they activate to ASCs in response to the infiltration of N. The presence of Md also contributes to the activation of QSCs represented by the rate *c*_*QSCMd*_ · *QSC* · *Md*. The process of QSC self-renewal by deactivation of ASCs has been hypothesized to depend on signals from M2 macrophages, which we model as *c*_*ASCM*2_ · *ASC* · *M* 2. In total, this produces the QSC dynamics equation:

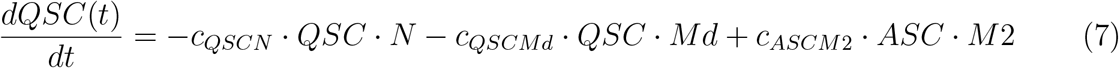

ASCs are initially converted from QSCs, depleting the number of QSCs. However, their subsequent proliferation is promoted by signals from proinflammatory M1 macrophages at the rate of 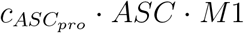. Conversely, M2 macrophages promote ASC differentiation, which we model as 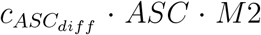. Finally, any ASC cells that neither proliferate, differentiation or return to quiescence, we assume somehow exit the system or are subject to apoptosis by the rate 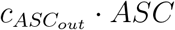.

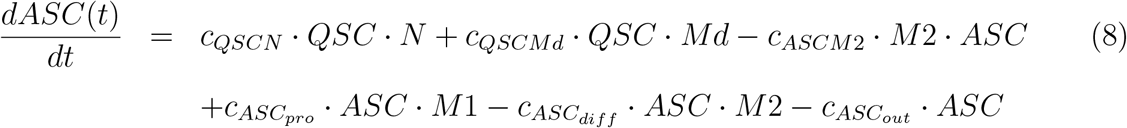

The equation for myoctyes is derived considering differentiation from ASCs and their “exit” from the system model, which typically happens when they fuse into multinucleated myofibers, which we do not model. The bilinear term 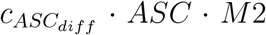 models the differentiation of ASCs into myocytes, while 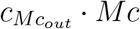 captures their exit from the model.

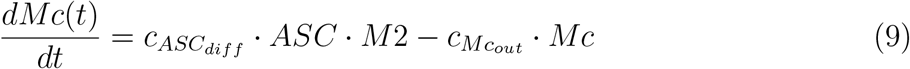

#### Initial conditions

To model an injury-regeneration dynamic, we assume that most cell types are not immediately present in significant numbers at the start of the simulation. Specifically, the initial conditions for all immune cells (N, Nd, M, M1, and M2), as well as downstream myogenic lineage cells (ASC and Mc) are set to zero. This assumption is based on the premise that these cells do not infiltrate the system immediately and are present in negligible numbers initially. Conversely, the initial condition for QSCs is set to 2700 cells/mm^3^, representing their baseline presence in the muscle tissue, and Md at approximately 30,000 cells/mm^3^, which reflects typical strong-injury scenarios as documented in the literature [19, 25, 3].

### 2.2 Establishing empirical cell type proportions time-series from scRNA-seq data

To establish a time-series of expected values for each cell type, we re-analyzed scRNA-seq data published by McKellar et al. [39]. This study looked at regeneration of the tibialis anterior (TA) muscle of mice after acute injury from notexin. The data included a total of 93,954 cells after quality control (QC), at six time points after injury: days 0, 1, 2, 3.5, 5 and 7 (Table 1). Applying the default Seurat pipeline we found 22 distinct cell type clusters (Fig. 2A), which we annotated by finding differentially expressed genes (DEGs) and marker genes for each cluster [18]. Among these are cell types such as MuSCs (see also Fig. 2B), immune cells (neutrophils and the monocyte-macrophage lineage), fibroadipogenic cells, endothelial cells, smooth muscle cells, pericytes, and tenocytes. Further validation of these cell types and their sub-clusters was conducted with cell type-specific gene expression profiles established in existing literature, demonstrated by the dot plots presented in Fig. 2C & 2D. For a comprehensive list of all the marker genes we analyzed, please refer to Supplemetary Table S1.

**Table 1:** Overview of scRNA-seq datasets used for modeling muscle regeneration.

| **Dataset** | **Age**<br>**(mo.)** | **# raw cell**<br>**counts** | **# cells**<br>**after QC** | **# samples** | **# time**<br>**points** | **Injury**<br>**days** |
| --- | --- | --- | --- | --- | --- | --- |
| McKellar et al. | 20 | 135,823 | 93,954 | 21 | 6 | 0, 1, 2, 3.5, 5, 7 |
| De Micheli et al. | 4-7 | 93,955 | 65,126 | 10 | 4 | 0, 2, 5, 7 |

**Figure 2.**
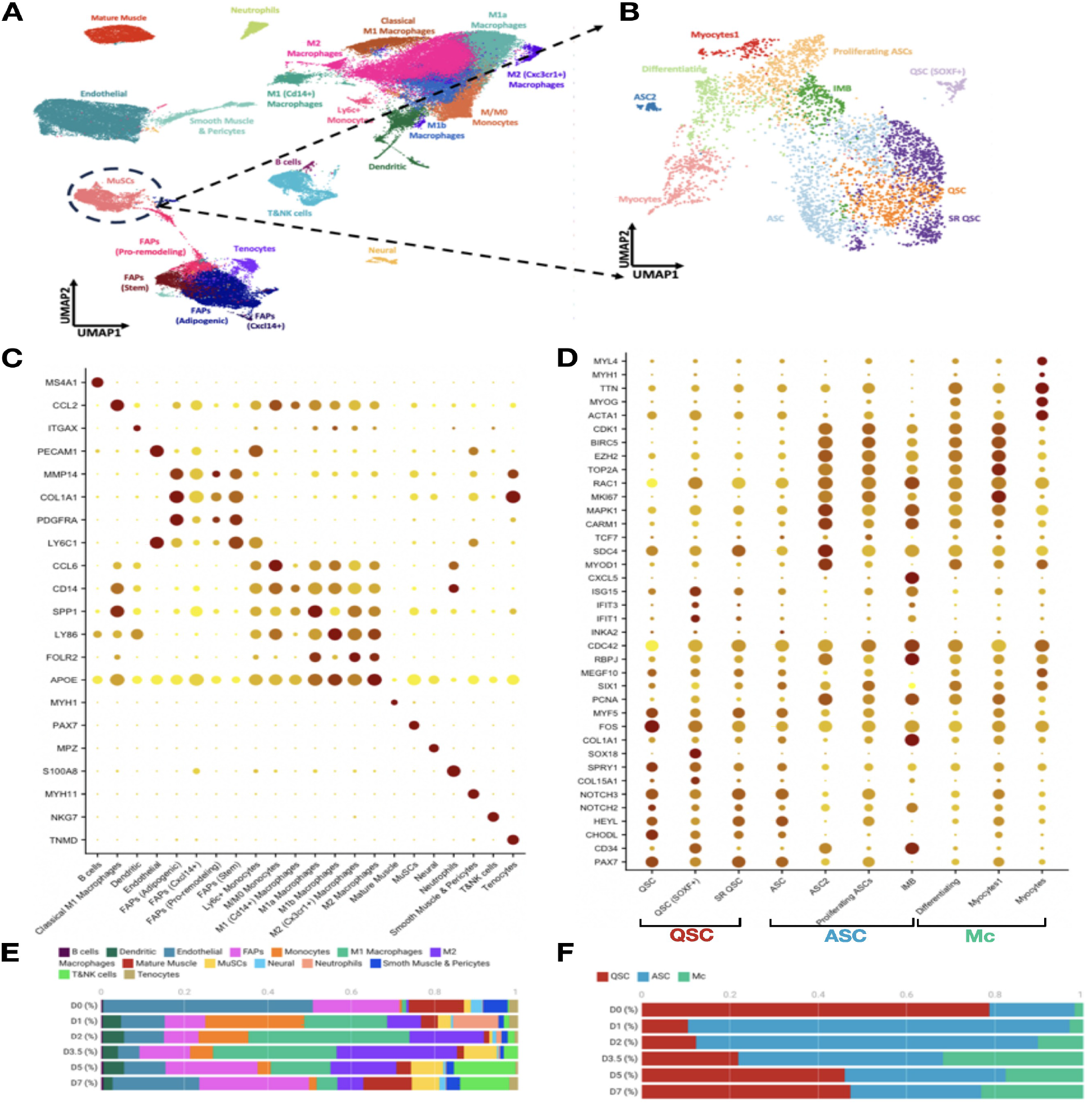
Comprehensive analysis of McKellar’s dataset revealing cell type dynamics during muscle regeneration. (A) UMAP projection of 93,954 single cells from integrated samples from the dataset by McKellar et al. Cells are color-coded based on cluster identity, illustrating the complex cellular ecosystem of the regenerative muscle tissue. Notable populations include various immune cell subsets, FAPs, and MuSCs, among others. (B) UMAP embedding of the MuSC subcluster displaying the temporal evolution of MuSC subtypes. Subtypes of MuSCs are differentiated by color, revealing the trajectory of MuSC activation, proliferation, and differentiation over the course of regeneration. (C) Dot plot validation of marker genes, where each dot’s size indicates the proportion of cells expressing that gene, and color intensity reflects the relative expression level (D) Dot plot validation of marker genes for MuSC sub-types identified in the literature. (E) Bar chart of cell count proportions calculated for each DPI with bars colored according to all cell type as shown in the UMAP projection in (A). (F) Bar chart of cell count proportions calculated for each DPI with bars colored according to MuSC sub-types as shown in the UMAP projection in (B).

From the cell-type-annotated scRNA-seq dataset, we calculated the proportion of each distinct cell type at each time point, as a fraction of the total cells at that time point (after aggregating cell subtypes, such as different types of M1 macrophages, QSCs and ASCs) (Fig. 2E, Table S2). These proportions were qualitatively consistent with biological expectations. For instance, at D0, immediately after injury, large numbers of endothelial and FAPs are present, but relatively few MuSCs, which only become significant after several rounds of proliferation. We also noted the presence of mature muscle at D0, even though healthy mature muscle cells are in the form of multi-nucleated myofibers that would not be expected to show up in a single-cell assay. This is likely due to experimental procedures including tissue mincing, digestion and filtering out of multinucleated myofibers during single cell isolation, which may result in some isolated or fragmented mature muscle-like cells [45, 7]. There was a transient increase in neutrophils at D1 (11%), rapidly declining thereafter, consistent with an immediate innate immune response post-injury. Monocytes surged to 24% at D1 and then peaked at D2 (39%), before gradually decreasing to 5% by D7. M2 macrophages showed a later peak at D3.5 (29%), which then diminished to 6% by D7, aligning with a transition from an inflammatory to a reparative tissue environment.

MuSCs consisted of 4,517 total cells, which we subclustered into quiescent, activated, and differentiating states, each manually annotated by distinct gene expressions (Fig 2D). Initially, the majority of satellite cells are in a quiescent state (QSCs: 79%), indicative of a typical resting muscle prior to (and immediately after) injury. It is likely that the trauma of injury at D0 rapidly induced a subset of QSCs to exit quiescence, thereby artificially lowering their apparent numbers when measured at the initial time point. Notably, the proportion of QSCs recovers from an initial drop (to 10%) back towards baseline levels (47% by the final time point), suggesting a restoration of the quiescent cell pool due to self-renewal processes. The relative stability in the proportion of QSCs towards the latter days may reflect a balance being struck between cell renewal and differentiation, ensuring sustained muscle regeneration capability. Upon injury, there is a dramatic shift as QSCs rapidly activate (ASCs peaking on day 1-2), entering the cell cycle to contribute to tissue repair. As the repair process progresses, the proportion of ASCs decreases (to 30% by the final time point), with an increase in the fraction of cells that have differentiated into mature myocytes (from 2% to 23%). This trend signifies the differentiation of ASCs to replenish the pool of functional muscle fibers as part of the healing process.

### 2.3 A second scRNA-seq dataset shows consistency of cell population proportions

The dataset by De Micheli et al. was selected to cross-validate both the numbers in the McKellar dataset and our model’s predictions [7]. This dataset contained 65,126 cells after QC from days 0, 2, 5, and 7 post-injury (Table 1). The data was put through the same analyses as the McKellar dataset, identifying cell types, producing counts of each cell type per time point, and transforming those into proportions (Table S3). Initially, monocytes comprised less than 1% of the cell population at D0, increased substantially to 10% by D2, and then decreased again to 1% by D7. Similar to the McKellar dataset, this pattern suggests an early influx of monocytes to the site of injury. The M1 and M2 macrophage populations show a pronounced increase at D2, with 33% and 28%, respectively, before decreasing.

In this dataset, a total of 2,478 MuSCs were identified. Of these, QSCs constituted 69% at D0, suggesting a predominantly quiescent state. However, this percentage decreased sharply to 4% by D2 and then partially recovered to 53% by D7. This pattern mirrors observations in McKellar’s dataset, where it is postulated that the initial injury and subsequent muscle dissociation processes at D0 might trigger a rapid transition of a subset of QSCs out of quiescence, leading to a temporary underrepresentation in their early counts. Indeed, the lower numbers of QSCs at D2 are consistent with their activation, and their rebounding by D7 is consistent with self-renewal. Following injury, the significant activation of satellite cells leads to ASCs peaking at 84% of MuSCs by D2. By D7, the ASC population diminishes to 33%, coinciding with a notable increase in differentiated myocytes (Mn).

The results from both datasets showed consistency in the proportions of major cell populations such as neutrophils, MuSCs, and some macrophage subsets across the different time points of muscle regeneration (Fig 3A,B, Tables S2 & S3). The immune cell dynamics within the McKellar dataset displayed a pronounced peak in neutrophil proportions (shown as a solid line in pink) at day 1 post-injury. Interestingly, the dataset by De Micheli et al. demonstrated a peak for neutrophils at day 2 (shown as a dashed line in pink), but the proportion was only half of that observed in the McKellar dataset at that day. This discrepancy could stem from the absence of day 1 data in De Micheli’s study, potentially obscuring the full neutrophil response. The trend for monocytes (shown as orange) followed a similar pattern. Consistency was more apparent in the proportions of M1 and M2 macrophages across both datasets, likely attributable to their sustained presence within the muscle regeneration milieu. MuSCs (Fig 3B, Tables S2 & S3) also exhibited comparable dynamics, with both datasets revealing a similar pattern of depletion and subsequent partial recovery, though neither returned to the original baseline levels observed at day 0. This could be attributed to technical artifacts from the experimental protocol that may have inadvertently activated these cells, or a naturally longer timeline required for the stem cell pool to replenish. ASCs (shown in blue) at day 2 presented comparable levels in both datasets. The lack of day 1 data in the De Micheli dataset (Fig 3B), however, precludes us from confirming the rapid activation of ASCs that was evident in McKellar’s data. For myocytes, the dataset by McKellar et al. showed a peak at day 3.5 post-injury. In contrast, the dataset by De Micheli et al. displayed a peak at day 5, nearly 25% lower than McKellar’s, suggesting that by day 5, a portion of myocytes might have already fused into new fibers, which is consistent with the muscle regeneration timeline. Based on these observations, it is plausible that the peak level of myocytes occurs somewhere between days 3 and 4 post-injury.

**Figure 3.**
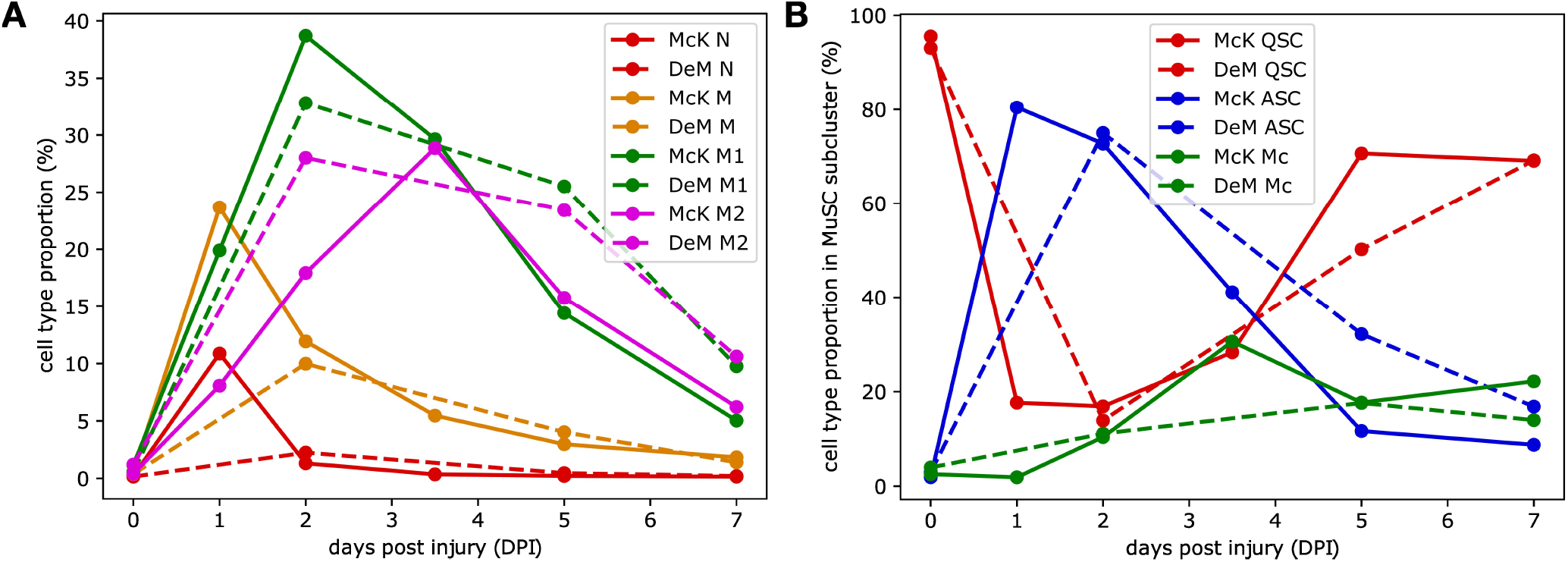
Cell-type population fractions from De Micheli’s and McKellar’s datasets are consistent. Line graphs plotting the proportions of key cell types across the time points for McKellar and De Micheli datasets show similar population dynamics in immune cells (A) and myogenic lineage cells (B).

### 2.4 Calibration of cell proportions to cell counts per cubic millimeter

Our mathematical model expresses cell-type abundances in units of cells per cubic millimeter, whereas annotated scRNA-seq data tells us only cell counts, or proportions, out of some indefinite volume or amount. By relying on published abundances of different cell types at certain point during regeneration, the proportions of the cells were calibrated to standardized units of cells per cubic millimeter (cells/mm^3^). Tables S2 and S3 contain the fractional proportions of all cells, including immune cells and MuSCs, relative to the total cell counts for each specific day post-injury from McKellar’s and De Micheli’s single-cell data. Table S4 shows literature values we found for different cell types at different days post injury, in similar injury models. Using those values, we determined scale factors such that if we multiply them by corresponding the scRNA-seq cell type percentages, we obtain the observed Table S2 or S3 cell count values. We applied the same scale factors to every cell type that we model, to obtain calibrated counts per day (see Table 2). For detailed calculations of the assigned scale factors at each DPI, please refer to the Methods and Materials section.

**Table 2:** Scaled cell counts per cubic millimeter, based on McKellar’s and De Micheli’s datasets, as a function of days post injury.

| Cell | McKellar |  |  |  |  |  | De Micheli |  |  |  |
| --- | --- | --- | --- | --- | --- | --- | --- | --- | --- | --- |
|  | D0 | D1 | D2 | D3.5 | D6 | D7 | D0 | D2 | D5 | D7 |
| N | 383 | 4500 | 273 | 134 | 98 | 77 | 426 | 548 | 140 | 59 |
| M | 950 | 9799 | 2466 | 2043 | 1324 | 887 | 1037 | 2433 | 1204 | 396 |
| M1 | 1967 | 8328 | 8000 | 11073 | 6400 | 2431 | 1436 | 8000 | 7624 | 2760 |
| M2 | 700 | 3341 | 3706 | 10781 | 7000 | 3000 | 3218 | 6830 | 7000 | 3000 |
| QSC | 2500 | 232 | 27 | 833 | 2447 | 2209 | 2500 | 24 | 989 | 1162 |
| ASC | 50 | 1056 | 116 | 1210 | 404 | 280 | 80 | 132 | 633 | 283 |
| Mc | 67 | 25 | 17 | 900 | 613 | 709 | 106 | 20 | 347 | 235 |

### 2.5 The ODE model can quantitatively capture regeneration dynamics

Initial values for all model parameters were first estimated by hand, using rough reasoning about time scales involved, approximate minimum and maximum expected values of different variables, and by-eye comparison of simulated model trajectories to the McKellar data–but not the De Micheli data. Then, the Nelder-Mead method was used to further optimize the model parameters, seeking to minimize total normalized absolute deviation between simulated and observed data. The initial and optimized parameters are listed in Table 3, while the optimized model simulations are shown in Fig 4. These simulations represent the predicted dynamics of cell populations in comparison with empirical single-cell data over a seven-day period following muscle injury.

**Table 3:** Initial and optimized parameter values for ODE model of muscle regeneration.

| Parameter | Description | Initial | Optimized | Units |
| --- | --- | --- | --- | --- |
| $c_{NMd}$ | Neutrophils engulfment of dead myonuclei | 2e-4 | 1.223e-4 | $\frac{mm^3}{cells \cdot d}$ |
| $c_{M1Md}$ | M1 macrophages engulfment of dead myonuclei | 3e-5 | 9.696e-5 | $\frac{mm^3}{cells \cdot d}$ |
| $c_{N_{in}}$ | Neutrophils infiltration | 10 | 7.053 | $d^{-1}$ |
| $c_{N_{out}}$ | Neutrophils exfiltration | 10 | 13.73 | $d^{-1}$ |
| $c_{M1Nd}$ | M1 macrophages engulfment of dead neutrophils | 1e-4 | 1.323e-4 | $\frac{mm^3}{cells \cdot d}$ |
| $c_{M_{in}}$ | Monocytes recruitment by neutrophils signaling | 4 | 3.741 | $d^{-1}$ |
| $c_{MM1}$ | Monocyte to M1 conversion | 5e-5 | 4.335e-5 | $\frac{mm^3}{cells \cdot d}$ |
| $c_{M_{out}}$ | Monocyte exfiltration | 1.2 | 1.229 | $d^{-1}$ |
| $c_{M1_{out}}$ | M1 macrophage exfiltration | 1e-10 | 2.3220e-8 | $d^{-1}$ |
| $c_{M1M2}$ | M1 to M2 conversion | 1000 | 1150 | $\frac{mm^3}{cells \cdot d}$ |
| $c_{M2_{inhib}}$ | M1 to M2 conversion inhibition | 1000 | 1844 | $\frac{mm^3}{cells \cdot d}$ |
| $c_{M2_{out}}$ | M2 exfiltration | 0.3 | 0.2646 | $d^{-1}$ |
| $c_{QSCN}$ | QSC activation to ASC related to N | 2e-4 | 1.260e-4 | $\frac{mm^3}{cells \cdot d}$ |
| $c_{QSCMd}$ | QSC activation to ASC related to Md | 2e-5 | 1.577e-5 | $\frac{mm^3}{cells \cdot d}$ |
| $c_{ASC M2}$ | ASC deactivation to QSC | 1e-4 | 1.355e-4 | $\frac{mm^3}{cells \cdot d}$ |
| $c_{ASC_{pro}}$ | ASC proliferation | 5e-6 | 8.778e-6 | $\frac{mm^3}{cells \cdot d}$ |
| $c_{ASC_{diff}}$ | ASC differentiation to myocytes | 5e-5 | 4.930e-5 | $\frac{mm^3}{cells \cdot d}$ |
| $c_{ASC_{out}}$ | ASC leaving the system | 1e-10 | 8.219e-12 | $d^{-1}$ |
| $c_{Mc_{out}}$ | Myocytes leaving the system | 0.2 | 0.1967 | $d^{-1}$ |

**Figure 4.**
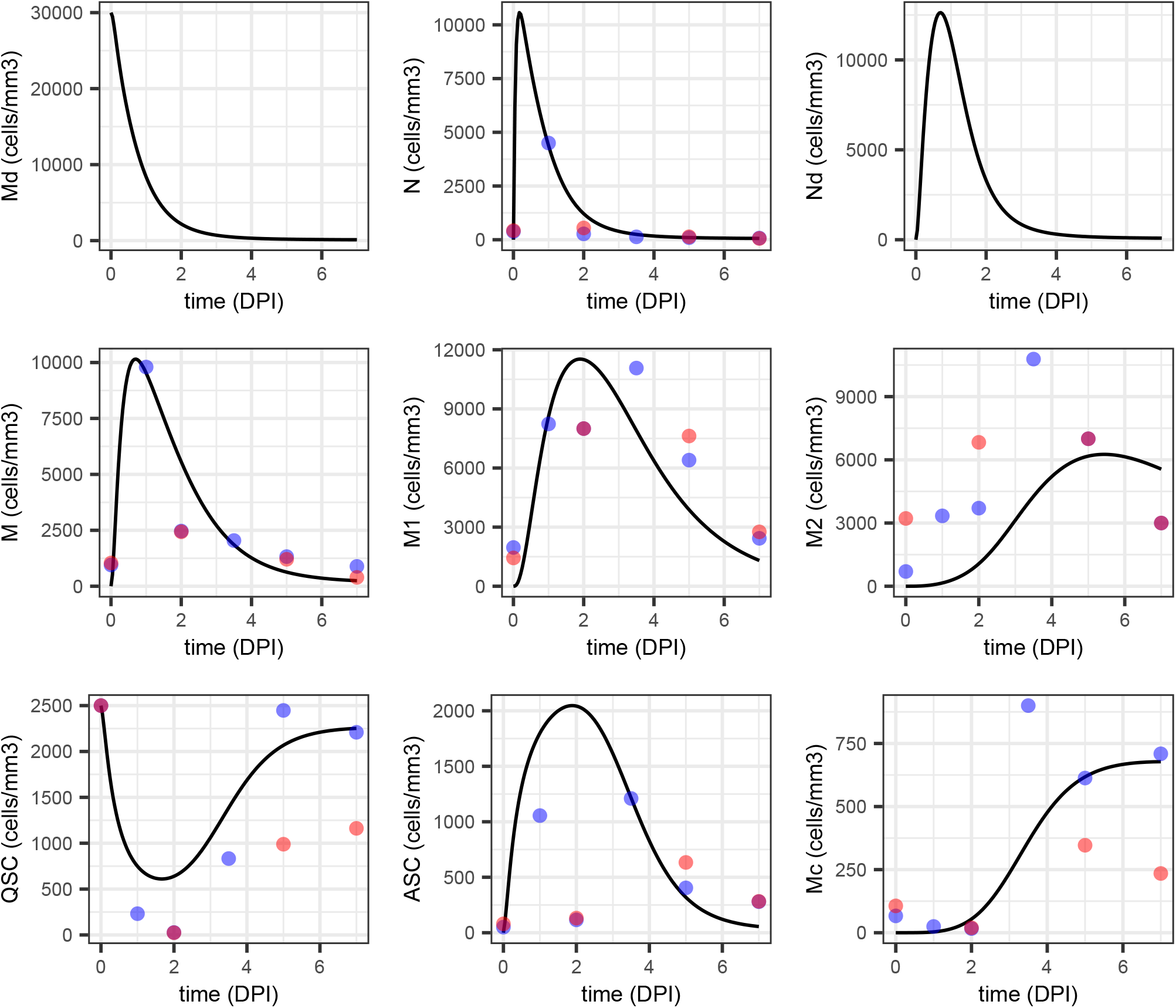
Model predictions versus empirical data over a seven-day time course following muscle injury. The solid black lines represent the trajectories obtained from the ODE model trained on the McKellar dataset. Empirical data points from the McKellar dataset are depicted as blue dots, while validation data from the De Micheli dataset are shown as red dots. Each subplot corresponds to a different cell population in the model: Md (dead myonuclei), N (neutrophils), Nd (dead neutrophils), M (monocytes), M1 (M1 macrophages), M2 (M2 macrophages), QSC (quiescent satellite cells), ASC (activated satellite cells), and Mc (myocytes). The empirical data do not include numbers for the dead cell types, Md and Nd. The alignment of model predictions with the McKellar (training dataset) data points indicates the model’s accuracy, whereas consistency with the De Micheli (validation dataset) data points supports the model’s generalizability.

Before optimization, our initial parameter estimates resulted in a simulated trajectory with 9.27 total error on the McKellar data, whereas the optimized trajectory has error 7.74. Detailed comparison of the simulated trajectories and empirical data can be found in the following paragraphs. However, simulation and data are in broad, qualitative agreement, and for some variables, in excellent quantitative agreement. The error on the De Micheli data, which was not part of the optimization procedure, was 17.5 with initial parameter estimates and 14.7 after optimization. This improvement in error on the held-out data suggests the model is not overfit and supports its generalizability.

Neutrophil counts were well-captured by the model for both datasets, but it is important to note that the model predicts an earlier peak for neutrophils than is visible in the data. In the simulated trajectory, neutrophil numbers peak at less than half a day into the time course. In the McKellar dataset, neutrophils peaks at day one, which is the earliest time point after the injury at day zero. The model captures that datapoint well, but predicts that neutrophil numbers are already declining rapidly, as they clean up dead myonuclei. The De Micheli dataset, which has its first post-injury time point at day 2, appears to miss the neutrophil peak, as by that time, both datasets show neutrophil numbers are comparatively small. Similarly, the simulation of monocytes also aligned closely with observed data, with monocyte numbers peaking in both simulation and empirical data at around one day post-injury.

The transition from M to M1 macrophages depicted by the model suggested a peak abundance around day two. In the McKellar data, the peak is at day 3.5, and in De Micheli at 2. Thus, the correct peak time is uncertain. But the peak magnitude shown by the model is approximately correct. The simulated M1 macrophage numbers drop a little earlier than the empirical data. Potentially, dead myonuclei Md, which are unobserved in the empirical data, persist longer than our model predicts, hence a need for M1 macrophages to engage longer in phagocytosis and signaling before differentiating into M2 macrophages.

The modeled M2 values are not an excellent fit to either empirical data set. Qualitatively, they follow a correct pattern of rising towards the latter half of the time series, when they send anti-inflammatory signals to the satellite cells, promoting differentiation into myocytes, to replace lost muscle, and return to quiescence, replenishing the dormant satellite cell pool. However, the M2 values appear to peak later than is observed in the data. Note, however, that there is considerable disagreement between the McKellar and De Micheli datasets on what M2 values should be. Also note that both empirical datasets suggest a considerable overlap in presence of M1 and M2 cell types, rather than a clean transition from one type to the other. This may be true globally in the muscle, or there may be local heterogeneity in how rapidly muscle regeneration proceeds, so that the scRNA-seq data is combining more cells from a more advanced regeneration state with cells in a less-advanced state.

The model trajectory of QSCs is qualitatively correct, dropping rapidly after injury, as the satellite cells activate, then rebounding close to original levels some days later, as a fraction of the proliferating satellite cells return to quiescence. The initial drop in QSC levels is not as strong as seen in either dataset. The rebound matches well the McKellar dataset, but less so the De Micheli dataset, which shows roughly half the numbers of QSCs as McKellar at days 5 and 7 post-injury.

The predicted ASC trajectory fits both datasets well, except at day 2, where the calibrated McKellar and De Micheli data both show quite low ASC numbers. The empirical numbers of ASCs at day 2 do not make qualitative sense, as we know ASCs are proliferating rapidly at that time. Possibly, the data is erroneous. The scaling factor resulting from our calibration may also be too small at that day. However, it cannot plausibly 10 or more times smaller than it should be, which is the amount that would be needed to bring the empirical ASC numbers into the ballpark of what would be expected based on the day 1 and 3.5 data. A 10 times larger scale factor for day 2 would make monocyte and macrophage numbers at that day implausibly large. Setting aside the issue with day 2, the qualitative shape of the ASC curve make sense, showing rapid expansion of ASC numbers due to proliferation following injury, succeeded by decline in their numbers as they either differentiate into myocytes or return to quiescence.

The model trajectory of myocytes, Mc, is similarly plausible, being very low initially, until satellite cells begin to differentiate late in the time series. McKellar and De Micheli datasets disagree by roughly a factor of two as to how many myocytes should be present late on. As described above, myocytes subsequently fuse to form myotubes, but we do not model that aspect of regeneration. Hence, in our model, the myocytes simply disappear from the model.

Collectively, these results show that the model is capable of producing qualitatively correct trajectories for all cell types, and for some cell types, the quantitative fit is quite good, matching both empirical datasets.

### 2.6 Sensitivity analysis reveals key model parameters

Sensitivity analysis was performed to quantify the responsiveness of model outputs to perturbations in each parameter. The derivative of each state variable at each time with respect to each parameter was estimated numerically based on a 1% perturbation in each parameter. The resulting derivatives are shown in Figure 5.

**Figure 5.**
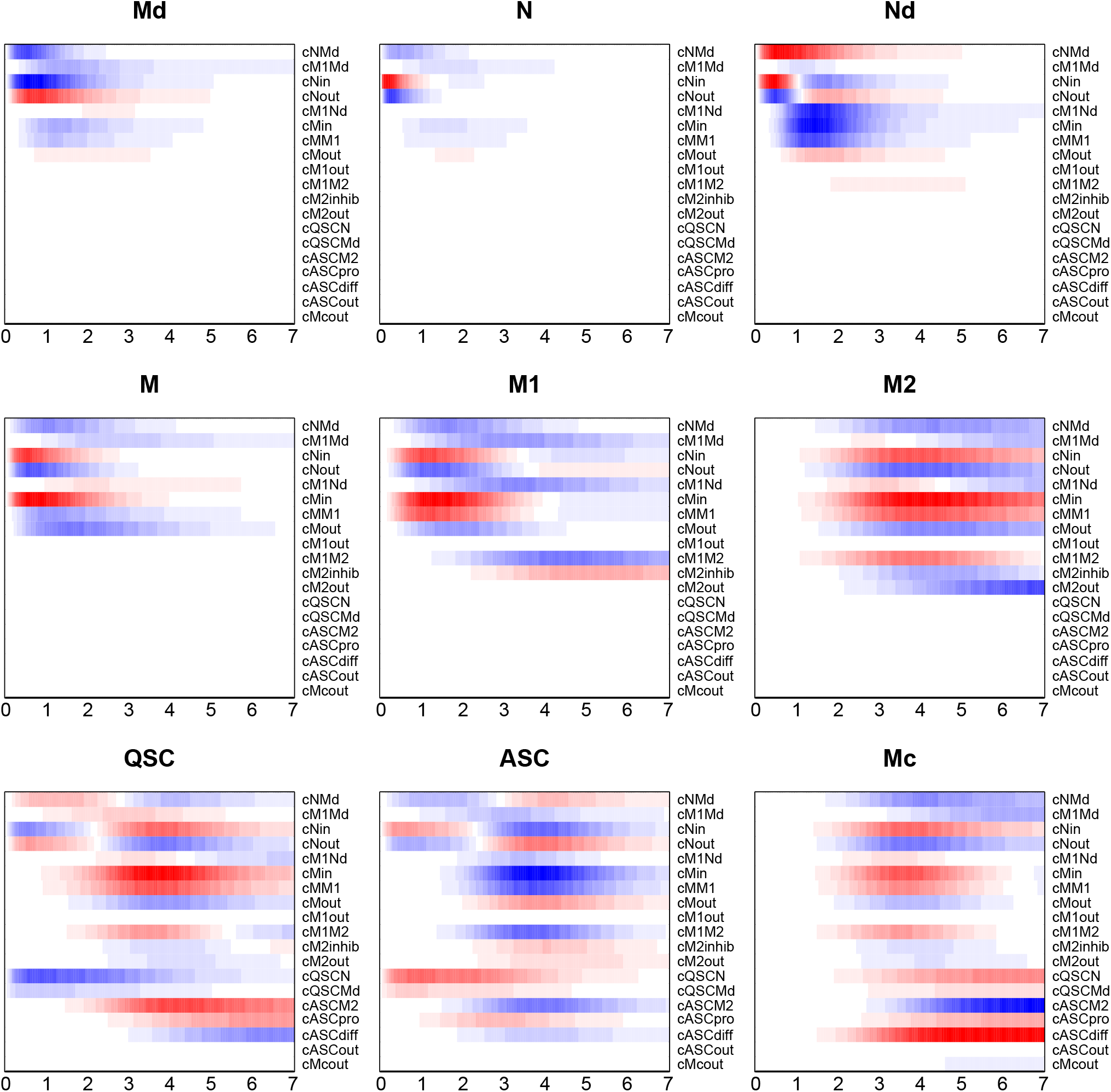
Sensitivity of modeled cell trajectories to parameter perturbations. Each heatmap shows the influence of a small positive perturbation to each model parameter on one of the nine state variables as a function of time. Red represents increase in the state variable, blue decrease. Within each heatmap, values are scaled to fill the full color range, so strengths of derivatives can be compared within a heatmap, but not across heatmaps.

For instance, the numbers of dead myonuclei Md between days 0 and 3 post injury are significantly reduced (blue) if *c*_*NMd*_, the rate at which neutrophils engulf dead myonuclei, is increased. Similarly, increasing the infiltration rate of neutrophils, 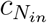, also causes a faster drop in dead myonuclei. Contrarily, increasing the rate at which neutrophils naturally exit the system, 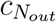, thereby reducing neutrophil numbers, results in more dead myonuclei remaining longer. The influences of monocyte and M1 macrophage parameters have similar effects on the dead myonuclei, but are lesser in magnitude, as the majority of the myonuclei clean up is due to neutrophils, according to the model. No parameter has a strong effect on dead myonuclei late in the time series, as by that time, they have been mostly cleared up and their numbers are near zero.

We do not examine every parameter’s influence on every state variable in detail, but we make several general observations and highlight some interesting cases. Notably, neither dead myonuclei nor any of the immune cells have any sensitivity to the satellite cell parameters. This is because, in our model, the influence goes strictly from immune cells to satellite cells, and there is no return influence. Second, most state variables are sensitive to multiple of the system parameters, often with several different parameters causing an increase and several others causing a decrease. In many cases, the directions of influence are obvious. For instance, increasing an infiltration or proliferation rate creates more of the corresponding cell, while increasing exfiltration or differentiation causes a decrease. However, some parameters have a bidirectional effect on a state variable, depending on which time during regeneration is considered. For instance, increasing the infiltration of neutrophils, 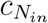, boosts the numbers of dead neutrophils during the first day, but reduces their numbers afterwards. The obvious reason for the increase is that if more neutrophils enter the system quickly, then more of them engulf dead myonuclei, and then themselves die, leading to increased dead neutrophils. However, the increase in early neutrophils also brings increased numbers of monocytes, which differentiate into M1 macrophages. Those can clean up dead neutrophils faster, leading to a drop in dead neutrophil numbers in the later days, relative to the nominal trajectory. Unsurprisingly, the neutrophil exfiltration rate, 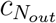, has the opposite effect on dead neutrophils, decreasing their numbers early on, but increasing their numbers later.

The monocyte-lineage cells are affected by numerous exfiltration and differentiation parameters, but a common thread to all of them, in addition to the neutrophil effects mentioned above, is the monocyte infiltration rate, 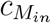, which tends to boost all monocyte-lineage cell numbers.

The satellite cells lineage is affected by most of the parameters, including of course the rate constants for the various activation, proliferation, differentiation, and deactivation processes. However, because those processes are also regulated by dead myonuclei numbers and different immune cells, the parameters that influence those cell types in turn influence the satellite cells as well. The sensitivities of the quiescent and activated satellite cells are nearly opposite. The main exception is the activated satellite cell proliferation rate, 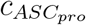, which increases both ASC numbers and QSC numbers, after the ASCs return to quiescence. Notably, one of the parameters with the strongest influence on QSC numbers is 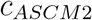, which controls the M2 macrophage influence on satellite cell deactivation. Our initial model did not include any such link, as there was not strong support in the literature. However, we found that this link, and proper tuning of its rate constant, was absolutely critical in ensuring the right number of activated satellite cells return to quiescence, and at the right time.

## 3 Discussion

In this article, we presented a population dynamic model that quantitively captures the dynamics of skeletal muscle regeneration following injury. Our model was trained on a single comprehensive scRNA-seq dataset, enhancing accuracy, and avoiding cell type misrepresentation, as subtle populations can influence the global dynamics. This dataset not only corroborated established findings, such as the successive presence of pro-inflammatory and anti-inflammatory macrophages during the regeneration cycle, but also enabled the classification of MuSCs with further manual annotation. Notably, the single-cell data reveals a predominantly quiescent satellite cell state at the onset, followed by a rapid transition to activated and committed states (Fig. 2F & Fig. 3B).

The model also makes several novel predictions about cell population dynamics and interactions. For one, the model predictions suggest that the influx of immune cells, specifically neutrophils and monocytes, may reach peak levels that are higher and/or earlier than those recorded in experimental data. Thus, it would be valuable to generate new experimental data looking in more detail at immune cell dynamics in the hours immediately post-injury. Because immune cell and satellite cell dynamics are orchestrated by complex interactions within the system, if these cell types are not fully captured by current experimental data, there may be important insights to gain that could lead to new hypotheses on cell signaling, cell fate decisions, and ultimately perhaps, new therapeutic avenues.

Another key finding relates to the effect of M2 macrophages on ASCs. The role of M2’s in promoting differentiation is very well established [1, 49], and while it has been suggested that they may also promote quiescence and self-renewal via contributions they make to the extra-cellular matrix [14], we are not aware of definitive evidence of this. We added this feature to our model when early testing without the term showed an inability of the model to both maintain ASC proliferation long enough and yet correctly replenish the QSC pool as proliferation winds down. Therefore, we advocate for further research to explore the role of M2 macrophages in the return of satellite cells to quiescence.

The proposed model enables, in principle, prediction of changes in cell population dynamics in response to molecular manipulations. For instance, one can modulate a single parameter–perhaps simulating up-or down-regulation of a key signaling factor between cell types–sand forecast shifts in the regeneration outcome or timeline. However, it is important to acknowledge that our model is a simplified representation, serving as a platform for future expansion and refinement. For instance, nearly all terms appear linearly, but nonlinear effects are widespread in cell biology. The model also incorporates no time delays that are inevitable when cells need to carry out fate decisions or migrate between compartments of the body. Still, having established that the model is able to account for wild-type regeneration dynamics, an important direction for future research is to test the model in perturbation conditions, and if necessary refine or expand the model to capture these conditions as well.

Despite the strengths of scRNA-seq in allowing us to quantify different cell types as a function of time, all datasets have their limitations. Our study employed two time-series datasets, one for training and one for validation of the model. The datasets largely agreed with each other, but for some cell types and at some time points they disagreed by a factor of two or three. Moreover, some data points, such as estimated ASC numbers at 2 days post injury, seem unlikely to be correct in either datasets. Another notable limitation is the inability of scRNA-seq to capture healthy myonuclei, as these exist in enormous, multi-nucleated myofibers. These can be captured by single-nucleus RNA-seq, but because they are so dominant in numbers, such data lacks precision for examining satellite cells. Thus, future research integrating single-cell and single-nuclei sequencing could provide a more accurate understanding of both damaged and regenerated muscle. Additionally, spatial transcriptomics, which allows for the mapping of gene expression in the context of tissue architecture, has enabled the identification of transcriptionally distinct nuclei originating from the same myofiber [4, 27, 24]. Therefore, the integration of single-cell sequencing, nuclei sequencing, and spatial transcriptomics in future muscle regeneration studies could significantly enhance our understanding of the full tissue microenvironment.

Despite these limitations, our model represents an import step towards improved understanding of cell population dynamics during regeneration. It includes more cell types and cell-cell interactions than any previous model. It correctly captures the main features of regeneration and explains them in terms of feedback rules between cell populations. In the future, we intend to expand the model to capture the misregulation or failure of regeneration in conditions such as Duchenne muscular dystrophy [13]. Additionally, we intend to include yet more cell types that influence satellite cells, most importantly, the fibro-adipogenic cells that proliferate alongside satellite cells after muscle damage, fill the muscle with fatty or fibrotic tissue, when satellite cells are unable to repair the muscle [6].

## 4 Materials and Methods

### 4.1 scRNA-seq data analysis

#### Data acquisition

Two publicly available datasets sourced from the tibilalis anterior (TA) muscles of C57BL6 mice were re-analyzed. For model training, we selected the dataset from McKellar et al. [39], containing 21 scRNA libraries from 20-month-old mice post-NTX injection. Validation was performed using the dataset by De Micheli et al. [7], which contained 10 libraries from 4–7-month-old mice at various time points after NTX-induced muscle damage. All libraries were originally constructed using the Chromium 3’ Library and Gel Bead Kits from 10X Genomics v3, USA and Illumina NextSeq 500 sequencing platform. The processed UMI count matrix files for the external datasets were acquired from GEO, which housed raw scRNA-seq data that have been pre-processed using Cell Ranger. These files correspond to datasets with the accession numbers SRP294168, and SRP241205, originally deposited in the Sequence Read Archive (SRA).

#### Data pre-processing and downstream analysis

Raw read counts provided by UMI count matrices for all samples were processed individually and combined among the other samples from the same source using the Seurat (v4) [18] within R v4.1.3 [57]. Samples were first loaded into R using the ‘Read10X’ function. Each sample’s gene expression matrix was then converted into a Seurat object using ‘CreateSeuratObject’. To filter out low-quality cells, genes that were expressed in fewer than 10 cells as well as cells with *<*1000 UMIs and *<*250 genes were removed from the gene expression matrix. In addition, we removed cells with *>*25% UMIs mapped to mitochondrial genes, as well as cell doublets with the ‘DoubletFinder’ function. The data was log-normalized using the ‘NormalizeData’ function in Seurat, setting a scaling factor of 10,000. Following this, gene expression was scaled by applying the ‘ScaleData’ function, which regressed out the effects of both the number of UMIs and the percentage of mitochondrial gene expression in each cell. Variable features across the datasets were identified using the ‘FindVariableFeatures’ function, setting the selection method to ‘vst’ with a feature count of 2,000. To address batch effects between samples, we employed the Harmony [30] integration method with default parameters. Visualization of the batch-corrected PCA indicated successful integration, with cells from different batches mixing well in the shared space. Following Harmony integration, dimensionality reduction was performed using PCA on the gene expression matrices and used the first 20 principal components (determined by ‘ElbowPlot’ function) for clustering and visualization using ‘RunPCA’, neighborhood graph construction using ‘FindNeighbors’, and clustering via ‘FindClusters’. Within the FindClusters algorithm, K-nearest neighbor (KNN) was calculated within Seurat to detect communities of cells. The ‘FindClusters’ function was executed with a resolution of 0.8, which was visualized and dimensionally reduced by the UMAP technique, using the ‘RunUMAP’ function. Finally, differential expression analysis was achieved using the ‘FindAllMarkers’ function using a likelihood ratio test that assumes the data follows a negative binomial distribution and only considering genes with *>*log2(0.25) fold-change and expressed in at least 25% of cells in the cluster.

#### Cell type annotation and sub-clustering

We applied a combined annotation approach for robust cell type labelling. This approach integrated both automated and manual annotation methods to capitalize on their respective strengths and mitigate their limitations. The initial step of the annotation process involved automated label transfer using a comprehensive reference atlas from a large-scale single cell/nuclei RNA-seq integration study [39]. Employing Seurat’s ‘FindTransferAnchors’ and ‘TransferData’ functions, the dataset was aligned to the reference atlas. This step enabled the assignment of cell type labels to our data based on transcriptional similarity with the predefined cell labels in the atlas. This reference dataset was solely utilized as an annotation tool, and its cells were not integrated into our primary analysis. Dimensionality reduction techniques, such as t-distributed Stochastic Neighbor Embedding (t-SNE) or UMAP, facilitated the visualization of cell clusters. Subsequent manual annotations were based on the expression patterns of well-established marker genes by referencing established literature for cell type identification. The final cell type labels were determined by comparing the outcomes of the automated and manual annotations. Upon completing the primary cell type annotation, we conducted high-resolution clustering on selected clusters for more in-depth analysis. For example, the MuSCs cluster was further divided using the ‘FindClusters’ function with a higher resolution setting. This detailed clustering was guided by established marker genes, enabling the categorization of MuSCs into different states such as quiescent, activated, and differentiating. These states were determined based on an extensive literature review.

### 4.2 Estimation and calibration of empirical time-series cell populations

The analyzed scRNA-seq data provided counts of different cell types identified at each day post injury. These counts offer a snapshot of the cellular composition at specific moments. To standardize these counts for comparison across time points, we converted them into proportions *P*_*cell*_ (Tables S2 & S3). This was achieved by dividing the count of each cell type by the total cell count at each time point. To translate the proportional data into absolute cell counts per mm^3^ of tissue, literature-reported cell densities for various cell types at specific DPIs were used as a reference. Scale factors were established for each reported day based on corresponding cell counts for one of the cell types from the literature (Table S4). These scale factors were determined for each DPI and applied to the proportions for all cell types.

Select studies report cell densities in terms of cells per milligram (mg) of tissue rather than cells per cubic millimeter, therefore their values were converted to a uniform volumetric density. This was accomplished by applying the established density of skeletal muscle tissue of 1.06 mg/mm^3^. Specifically, cell counts from Tonkin et al. [60] regarding macrophage populations, initially provided in cells/mg, were recalculated into cells/mm^3^ using the formula:

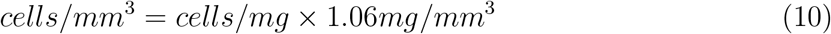

Similarly, for two-dimensional measurements such as cells per square millimeter (cells/mm^2^), we converted these to cells/mm^3^ by considering the standard histological section thickness of 10 *μ* m = 0.01 mm, as

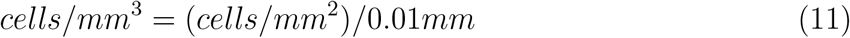

### 4.3 Parameter estimation

Parameters were estimated using a two-step process involving an initial approximation followed by numerical optimization. The initial parameter values were set based on a combination of approximate theoretical expectations, based on timing of certain cell populations’ increases and decreases, and eye-ball fit to the McKellar empirical data [39]. The Nelder-Mead algorithm, part of the R ‘optim’ function was used to further optimize parameters to fit the McKellar data, seeking to minimize the normalized, summed absolute error between modeled trajectory and empirical data. With observed cell counts *y*_*j*_(*t*_*i*_) for cell type *j*, and model-predicted cell counts *x*_*j*_(*t*_*i*_, *θ*), where *t*_*i*_ represents observations times and where *θ* is a vector of parameters, this error is:

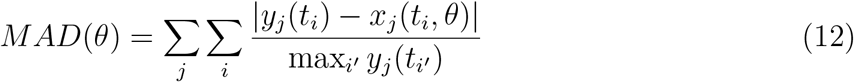

where *i* and *i*^*!*^ range over the observation times and *j* ranges over the cell types. The objective is to find the parameter vector *θ* that minimizes the error subject to the parameters *θ* being non-negative. To avoid the non-negativity constraint, we optimized unconstrained log-transformed parameters *θ*^*!*^ where *θ* = *ln*(*θ*^*!*^). For any given set of parameters, the modeled cell counts were obtained by solving the system of differential equations via the ‘ode’ function in R, using the ‘lsoda’ method. The initial conditions for the system of ODEs was Md = 30000, QSC = 2500, and all other state variables set to zero.

## 5 Funding

This work was supported in part by a grant from the Artificial Intelligence for Design (AI4D) challenge program from the National Research Council of Canada, and by grant RGPIN/06604-2019 from the Natural Sciences and Engineering Research Council of Canada.

## 6 Acknowledgements

We thank Vahab Soleimani, Jeffrey Dilworth, and Michael Rudnicki for many useful discussions about muscle regeneration, our model, and biological interpretation of the scRNA-seq data.

## Notes

### Competing Interest Statement

The authors have declared no competing interest.

