## Supplementary Information for "Differential Equation Modeling of Cell Population Dynamics in Skeletal Muscle Regeneration from Single-Cell Transcriptomic Data"

**Table S1.** Gene markers used for cell-type annotation.

| Cell Type | Marker Genes | Reference |
|---|---|---|
| **Endothelial** | PECAM1 (CD31), CDH5, VEGFR2 (KDR) | Imoukhuede & Popel, 2012; Kalucka et al., 2020 |
| **Smooth** | ACTA2, MYL9, MYH11 | Muhl et al., 2022 |
| **Pericytes** | MCAM, PDGFRB, NG2 (CSPG4) | Rippe et al., 2021 |
| **Tenocytes** | TNMD, SCX, COL1A1 | Gumucio et al., 2020 |
| **Mature** | MYH1, MYH2, MYH4, ACTA1 | Oprescu et al., 2020b; Williams et al., 2022; McKellar et al., 2021 |
| **FAPs (Adipogenic)** | PPARG, CD36 | Molina et al., 2021 |
| **FAPs (Stem)** | PDGFRα, SCA1 | McKellar et al., 2021 |
| **FAPs (Pro-remodeling)** | TGFβ, MMP14 | McKellar et al., 2021 |
| **FAPs (Cxc14+)** | CXCL14 | Oprescu et al., 2020b |
| **Neutrophils** | S100A8, S100A9, CXCR2, CCL3 | McKellar et al., 2021; Reichel et al., 2009 |
| **M0/M Monocytes** | CTSA, CTSL, CTSB, CCR2, CD14, ITGAM, LYZ2, S100A4, CCL2 | De Micheli et al., 2020; Ingersoll et al., 2010; McKellar et al., 2021; Oprescu et al., 2020b |
| **Ly6C+ Monocytes** | CD177, LY61C, LY6C2C, IRF5 | Corbin et al., 2020; Stroncek et al., 2004; Yang et al., 2021 |
| **Classical M1 Macrophages** | CLL2, CCL6, CCL9, TNF, IL10, NOS2 | Krasniewski et al., 2022a; McKellar et al., 2021; Corbin et al., 2020; Krasniewski et al., 2022b |
| **M1a Macrophages** | SPP1, IL17RA, CXCL3, CXCL16 | Mao et al., 2022; Patsalos et al., 2022; Zhang et al., 2009 |
| **M1b Macrophages** | CCR2, CXCL16, C1QC, IL6RA, CSF1R, AIF1 | De Micheli et al., 2020; Krasniewski et al., 2022a |
| **CD14+ Macrophages** | CD14, SLFN4, CXCL2, FOSL2, ATF4, MDM2 | De Micheli et al., 2020; Krasniewski et al., 2022a; McKellar et al., 2021; Oprescu et al., 2020b |
| **CX3CR1+ Macrophages** | CX3CR1, LY86, LPXN, DHRS3, DNMT3a | Wang et al., 2016 |
| **M2 Macrophages** | ARG1, CD163, MRC1, CCL8, FOLR2, APOE, TREM2 | De Micheli et al., 2020; Krasniewski et al., 2022a; McKellar et al., 2021; Oprescu et al., 2020b |
| **T cells** | CD3E, CD4, CCL5, CD8A | McKellar et al., 2021 |
| **NK cells** | NKG7, KLRA7, NCAM1 (CD56) | McKellar et al., 2021 |
| **B cells** | CD19, MS4A1 (CD20) | McKellar et al., 2021 |
| **Dendritic cells** | CD1C, HLA-DR,CD11c (ITGAX) | McKellar et al., 2021 |
| **QSCs** | Chodl, HeyL, Pax7, Sdc3 | Fukada et al., 2007, Barutcu 2022 |
| **Self-Renewing QSCs** | Notch2/3, Dpt, Calcr, Col15a1, Spry1 | Oprescu et al., 2020 |
| **ASCs (early activated)** | Myod1, Sdc4, Tcf7, Carm1, Mapk, Fos, Mest | Segalés et al., 2016, Jones et al., 2005, Le Grand et al., 2009, Barutcu 2022, De Micheli 2021 |
| **IMB** | Ctsb, Ifit1, Ifit3, Isg15, Cxcl | Oprescu et al., 2020 |
| **Proliferating ASCs** | Ki67, Rac1, Top2a, Ezh2, Birc5, Cdk1 | Oprescu et al., 2020 |
| **Myocytes & Differentiating** | Acta1, Myog, Ttn, Myh, Myl4 | McKellar et al., 2021 |

**Table S2.** Proportions (%) of Total Cells Across Time Points (0,1,2,3.5,5,7) in McKellar's Dataset

| Cell Type | D0 (%) | D1 (%) | D2 (%) | D3.5 (%) | D5 (%) | D7 (%) |
|---|---|---|---|---|---|---|
| **B cells** | 0.25% | 0.28% | 0.32% | 0.23% | 0.45% | 0.40% |
| **Dendritic** | 0.21% | 4.49% | 5.22% | 3.79% | 5.08% | 2.34% |
| **Endothelial** | 50.15% | 10.42% | 9.47% | 5.16% | 9.89% | 20.65% |
| **FAPs** | 20.82% | 9.86% | 8.23% | 12.13% | 22.14% | 26.32% |
| **Monocytes** | 0.57% | 23.67% | 11.93% | 5.47% | 2.98% | 1.84% |
| **M1 Macrophages** | 1.18% | 19.90% | 38.70% | 29.65% | 14.41% | 5.04% |
| **M2 Macrophages** | 0.42% | 8.07% | 17.93% | 28.87% | 15.76% | 6.22% |
| **Mature Muscle** | 13.42% | 3.94% | 1.22% | 1.70% | 3.49% | 11.62% |
| **QSCs** | 1.500% | 0.560% | 0.130% | 2.230% | 5.510% | 4.580% |
| **ASCs** | 0.030% | 2.550% | 0.560% | 3.240% | 0.910% | 0.580% |
| **Myocytes** | 0.040% | 0.060% | 0.080% | 2.410% | 1.380% | 1.470% |
| **Neural** | 2.77% | 0.61% | 0.93% | 0.34% | 0.55% | 1.47% |
| **Neutrophils** | 0.23% | 10.87% | 1.32% | 0.36% | 0.22% | 0.16% |
| **Smooth Muscle & Pericytes** | 5.76% | 1.23% | 1.39% | 0.94% | 1.84% | 3.42% |
| **T&NK cells** | 0.40% | 1.33% | 1.69% | 3.22% | 14.75% | 11.67% |
| **Tenocytes** | 2.23% | 2.16% | 0.90% | 0.27% | 0.65% | 2.23% |
| **Total** | 100.00% | 100.00% | 100.00% | 100.00% | 100.00% | 100.00% |

**Table S3.** Proportions (%) of Total Cells Across Time Points (0,2,5,7) in De Micheli's Dataset.

| Cell Type | D0 | D2 | D5 | D7 |
|---|---|---|---|---|
| **FAPs** | 52.57% | 9.27% | 18.74% | 35.60% |
| **B cells** | 0.17% | 0.97% | 0.54% | 0.74% |
| **Dendritic** | 0.86% | 6.43% | 4.90% | 3.21% |
| **Endothelial** | 31.65% | 5.07% | 4.77% | 17.24% |
| **Monocytes** | 0.39% | 9.98% | 4.03% | 1.40% |
| **M1 Macrophages** | 0.54% | 32.82% | 25.52% | 9.76% |
| **M2 Macrophages** | 1.21% | 28.02% | 23.43% | 10.61% |
| **Mature Muscle** | 3.53% | 0.25% | 0.80% | 1.48% |
| **QSC** | 0.94% | 0.10% | 3.31% | 4.11% |
| **ASC** | 0.03% | 0.54% | 2.12% | 1.00% |
| **Myocytes** | 0.04% | 0.08% | 1.16% | 0.83% |
| **Neural** | 1.79% | 0.16% | 0.24% | 1.41% |
| **Neutrophils** | 0.16% | 2.25% | 0.47% | 0.21% |
| **Smooth Muscle & Pericytes** | 3.43% | 0.58% | 0.71% | 2.89% |
| **T&NK cells** | 1.05% | 3.36% | 8.97% | 6.70% |
| **Tenocytes** | 1.62% | 0.13% | 0.31% | 2.81% |
| **Total** | 100.00% | 100.00% | 100.00% | 100.00% |

**Table S4.** Literature-Based Cell Counts.

| DPI | Study Reference | Cell Type | Cell Count/Measurement | Unit |
|---|---|---|---|---|
| 0 | Bouredji 2021, Keefe 2015, Hardy 2016 | QSCs | 2000, 2500, 2700 (median 2500) | cells/mm$^3$ |
| 1 | Dumont 2010, Pizza 2005 | Neutrophils | 3500-5500 (median 4500) | cells/mm$^3$ |
| 2 | Tidball 2007 | M1 Macrophages | 8000 | cells/mm$^3$ |
| 3.5 | Tidball 2007 | Myocytes | 900 | cells/mm$^3$ |
| 5 | Tonkin 2015 | M2 Macrophages | 7000 | cells/mg |
| 7 | Bouredji 2021 | M2 Macrophages | 3000 | cells/mm$^3$ |
